# phastSim: efficient simulation of sequence evolution for pandemic-scale datasets

**DOI:** 10.1101/2021.03.15.435416

**Authors:** Nicola De Maio, William Boulton, Lukas Weilguny, Conor R. Walker, Yatish Turakhia, Russell Corbett-Detig, Nick Goldman

**Author notes:** Corresponding author; Content. These authors contributed equally to this work (Nicola De Maio, William Boulton).

## Abstract

Sequence simulators are fundamental tools in bioinformatics, as they allow us to test data processing and inference tools, as well as being part of some inference methods. The ongoing surge in available sequence data is however testing the limits of our bioinformatics software. One example is the large number of SARS-CoV-2 genomes available, which are beyond the processing power of many methods, and simulating such large datasets is also proving difficult. Here we present a new algorithm and software for efficiently simulating sequence evolution along extremely large trees (e.g. < 100, 000 tips) when the branches of the tree are short, as is typical in genomic epidemiology. Our algorithm is based on the Gillespie approach, and implements an efficient multi-layered search tree structure that provides high computational efficiency by taking advantage of the fact that only a small proportion of the genome is likely to mutate at each branch of the considered phylogeny. Our open source software is available from https://github.com/NicolaDM/phastSim and allows easy integration with other Python packages as well as a variety of evolutionary models, including indel models and new hypermutatability models that we developed to more realistically represent SARS-CoV-2 genome evolution.

**Author summary:** One of the most influential responses to the SARS-CoV-2 pandemic has been the widespread adoption of genome sequencing to keep track of viral spread and evolution. This has resulted in vast availability of genomic sequence data, that, while extremely useful and promising, is also increasingly hard to store and process efficiently. An important task in the processing of this genetic data is simulation, that is, recreating potential histories of past and future virus evolution, to benchmark data analysis methods and make statistical inference. Here, we address the problem of efficiently simulating large numbers of closely related genomes, similar to those sequenced during SARS-CoV-2 pandemic, or indeed to most scenarios in genomic epidemiology. We develop a new algorithm to perform this task, that provides not only computational efficiency, but also extreme flexibility in terms of possible evolutionary models, allowing variation in mutation rates, non-stationary evolution, and indels; all phenomena that play an important role in SARS-CoV-2 evolution, as well as many other real-life epidemiological scenarios.

## Introduction

Sequence evolution simulation is important for many aspects of bioinformatics [1]. Its most ubiquitous applications are for testing and comparing the performance of various essential tools (such as alignment, phylogenetic, and molecular evolution inference software, see e.g. [2–4]). However, simulating sequence evolution is also often used for testing hypotheses (e.g. [5]) and for inference, either for example through Approximate Bayesian Computation [6, 7], see e.g. [8, 9], or, more recently, using deep learning, see e.g. [10–12].

Many simulators address the task of simulating gene trees, or ancestral recombination graphs, as well as simulating evolution along these trees (e.g. [13–16]). Instead, here we focus on the problem of generating sequences given an input tree, as done by “phylogenetic” simulators (e.g. [17–19]). Realistic simulation of sequence evolution along a phylogenetic tree is essential, for example, for assessing and improving our methods for inference of SARS-CoV-2 phylogenies, which is a largely still open problem [20]. One important factor is the large numbers of available genome sequences for SARS-CoV-2 (> 3, 000, 000 in the GISAID database [21] as of September 2021). Despite this, there are currently no available simulation frameworks capable of simulating the scale and complex evolutionary features of SARS-CoV-2 and similar genome datasets. For this reason, we focus on the issue of simulating realistic substitution patterns for large datasets of closely related samples, as broadly observed in genomic epidemiology sequence data, and for arbitrarily complex substitution and indel models.

Here we show that sequence simulation for such large numbers of genomes is exceedingly computationally demanding for existing software. Complex evolutionary models, for example codon substitution models and rate variation, can cause significant further slow-downs. Furthermore, many existing methods do not allow the simulation of mutational patterns realistic for SARS-CoV-2, such as non-stationary and highly variable mutational processes [22–24], or don’t allow the simulation of indels. We propose a new approach to efficiently simulate the evolution of many closely related genomes along a phylogenetic tree and under general sequence evolution models. Our approach simulates one mutation (substitution or indel) at a time using the Gillespie method [25], and is further tailored to reduce time and memory demand by efficiently representing and storing information regarding non-mutated positions of the genome. Furthermore, we use a multi-layered search tree structure to efficiently sample mutation events along the genome even when each position has its own mutation rate, and to efficiently traverse the phylogenetic tree and avoid redundant operations. Our approach empowers extremely flexible and fine-grained evolutionary models. For example, non-stationary models are specifiable, with each nucleotide position of the genome assigned a distinct mutational profile, and each codon a distinct nonsynonymous/synonymous rate ratio. Similarly flexible indel models are also specifiable.

## Materials and methods

We consider the problem of simulating evolution of a DNA (or RNA) sequence along a specified input phylogenetic tree, and under a given evolutionary model. Our simulation approach is based on the Gillespie method [25], as is typically used in molecular evolution simulators [18, 19]. We assume that each position of the genome (either nucleotide or codon) evolves independently of the others, and under a time-homogeneous substitution process; that is, the rates of evolution at each position are initially specified by the user or are sampled randomly by the simulator. We focus on the efficient simulation of sequence evolution for large phylogenetic trees with short branches: we assume that only a few mutations happen on each branch across the genome, which is typical for genomic epidemiology, and in particular for SARS-CoV-2 [26].

### “Vanilla” approach

If we assume that evolutionary rates are homogeneous across the genome, it is simple to use the Gillespie approach efficiently in this scenario by adopting an efficient representation of ancestral genomes in terms of differences with respect to a root genome [27]. As a very simplified example, let’s consider the case in which there is no selective force at play, mutation rates are constant across the genome, there are no indels, and all bases mutate into all other bases at the same rate (JC69 model [28] with equal nucleotide frequencies). Throughout the manuscript, we will not assume equilibrium or stationarity in sequence evolution, but instead assume that we are given a genome at the root of the phylogeny, which we then evolve down the tree according to given rates.

In this simplified “vanilla” scenario, the total mutation rate across the genome is equal to the mutation rate for one base, 3*r*, times the genome length (which we assume constant), *L*. Starting from the root and its genome, we visit each branch of the tree one at the time in preorder traversal. For each branch of the tree, we consider its length *t*_*b*_, and we recursively sample a time for the next mutation from an exponential distribution with parameter 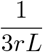. If the sampled time *t* is over *t*_*b*_, we move to the next branch. Otherwise, we decrease *t*_*b*_ by *t* and we sample a mutation event. In the considered scenario, this simply means sampling one position of the genome at random (a random integer number 1 ≤ *i* ≤ *L*), and then a random allele *b*, different from the current allele at position *i*, to mutate into. Additional steps are also required to keep track of mutations which have already occurred and allow them to further mutate, for example, possibly reversing a mutated allele back to the reference allele. We track each sampled mutation by adding it to a list of mutations for the current branch. It is worth noting however that there are more efficient ways to keep track of mutations that have already occurred, which we discuss in subsequent sections. A pseudocode description of the algorithm is given in Algorithm 1. So overall, the total cost of this “vanilla” algorithm is constant in genome size, and is linear in the number of tips *N*. It does however scale with the number of mutation events (total tree length) *M* = *O*(*Nl*) where *l* is the average number of mutations per branch. The initialization step has cost *O*(*N*) in order to read the phylogenetic tree, and further *O*(*L*) with more complex models in order to keep track of the positions of different alleles. Performing the simulations has cost *O*(*M* log(*N*) + *M*^2^ log(*N*)/*N*) = *O*(*l*^2^*N* log(*N*)); the main cost here is to screen previous mutation events at each new mutation, and this can be significantly reduced as explained in the next section. There is a caveat however. The default output of our software phastSim is a text file where each sample name is followed by a list of differences of the simulated sample genome with respect to the reference. We also allow users to print a Newick format tree, annotated with the simulated mutation events. If, however, we want to produce a file containing the full alignment, the memory and time cost of the algorithm will become *O*(*NL*), since this is the size of the alignment. For this reason, we provide the option for the user to generate a FASTA or PHYLIP alignment output, but by default we only generate the more concise version consisting of a list of differences, which usually leads to a very considerable reduction in time and memory demand.

#### Algorithm 1

Vanilla algorithm for one phylogenetic branch.

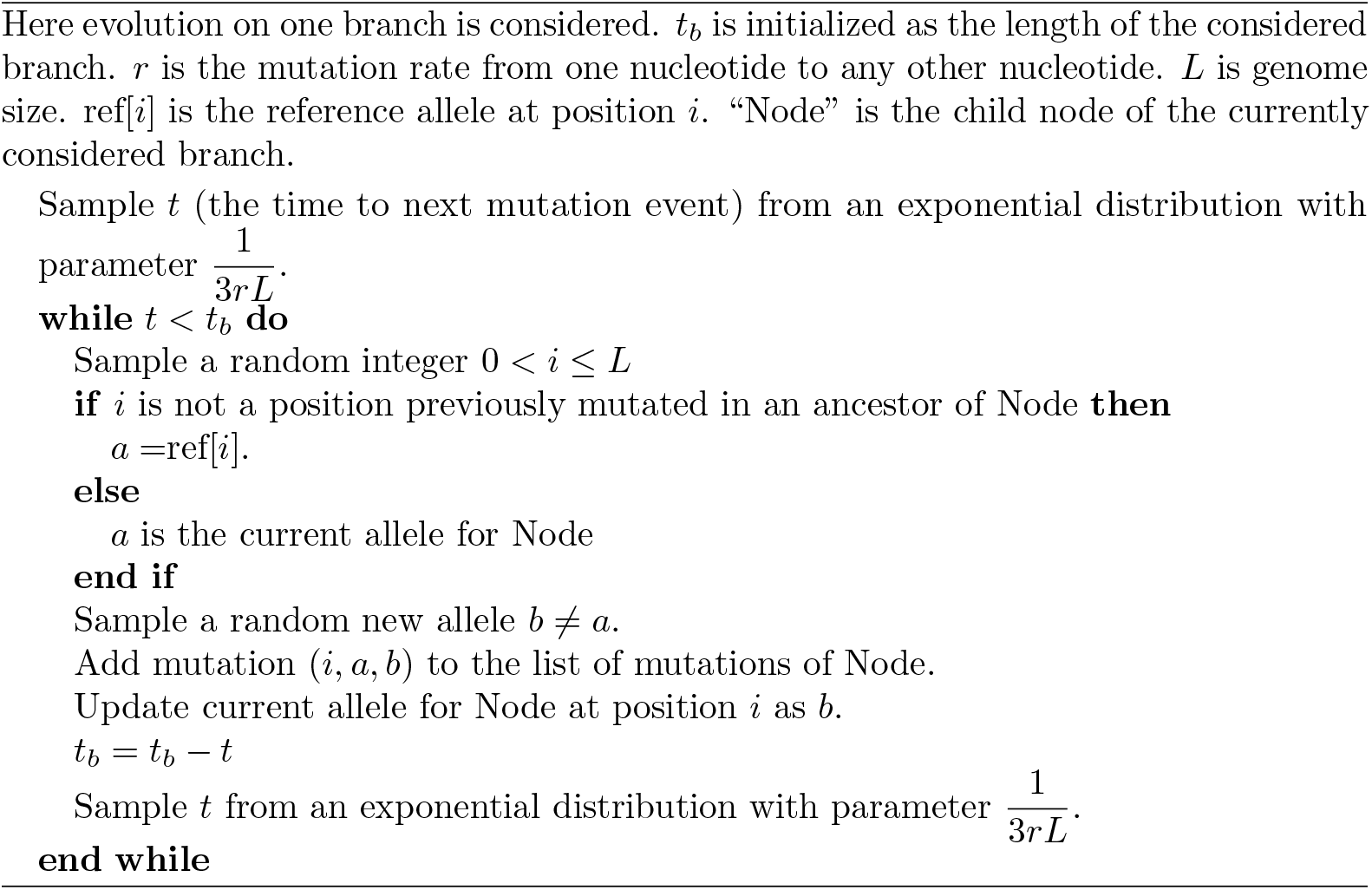

In classical implementations of sequence evolution simulators [17], for each node of the tree we need to update each base of the genome one at the time, therefore incurring in cost *O*(*NL*). Therefore, when the number of expected mutations is *M* ≪ *NL* we expect an advantage in using this approach.

A considerable limitation of the above “vanilla” approach is that we assume that rates are the same across the genome, and this is hardly realistic [29, 30]. We implemented a simple extension of this algorithm above which accounts for both an arbitrary nucleotide substitution model (UNREST [31]), and for rate variation across the genome in terms of a finite number of rate categories. To achieve this, we extended the algorithm above to keep track of which positions of the genome have which rates. This allows us to efficiently calculate total mutation rates for each class of sites, and to efficiently sample sites within a class.

We also implemented a new model of rate variation in order to better fit the patterns of hypermutability observed in SARS-CoV-2. In this model, small proportions of hypermutable sites are given a (possibly much) higher mutation rate. At an hypermutable site, only one specific mutation rate (from one nucleotide to one other nucleotide) is enhanced. For example, one such site with hypermutability might have only the G → T mutation rate increased 100-fold, while all other rates at that site remain the same. This is to model the effects observed in SARS-CoV-2 which are possibly attributable to APOBEC and ROS activity (or other still unclear mechanisms) [22, 23].

However, as the number of site classes increases, and as the number of alleles increase (for example when considering codon models), the efficiency of the extension of the vanilla approach described above deteriorates, especially when each site of the genome is given different evolutionary rates. For this reason, we developed a more complex algorithm that remains efficient in light of rate variation, with only a small efficiency sacrifice relative to the vanilla method in the scenario of no rate variation. We allow phastSim users to choose between the vanilla approach or the more complex one, that we call “hierarchical” and describe below. Advanced features, for example simulation of indels, are only implemented with the hierarchical algorithm.

### “Hierarchical” approach

#### Binary search “genome” tree

We first describe the structure and algorithm that allow us to efficiently sample a mutation event along the genome when each position might have a distinct mutation rate. This structure needs to be efficiently updatable following a mutation event; in fact, a mutation event changes the allele at a position of the genome, and therefore also its mutation rate. This is very similar to the problem of sampling from a categorical distribution with many elements, where the probabilities can be slightly modified at each sample [32]. A Huffman tree [33] would be close to optimal for this task, however, here we implement a binary search tree, which has a slightly higher expected cost [32] but allows us to more efficiently model blocks of contiguous nucleotides, and therefore to efficiently simulate indels.

In our “genome” search tree (which is distinct from, and should not be confused with, the phylogenetic tree), each node corresponds to a contiguous block of nucleotides along the genome. The root node represents the whole genome, and contains a rate value corresponding to the global mutation rate of the whole genome. The two children of the root correspond to the first and the second half of the genome, respectively. There is no overlap between the regions considered by each child node, and their union gives the region considered by the parent node. Consequently, the sum of the rates of the children of a node is equal to the rate of the node. Given this structure, we also refer to this binary search tree as the “genome” tree. A terminal node of the genome tree corresponds to one unit of the genome, either a base or a codon, depending on the model we choose for simulations. A terminal node contains not only information about the position of the unit along the genome, but also the reference allele at this position and the mutation rates associated with it, to allow sampling of a specific mutation event at the given position/node. A graphical representation of an example genome tree is depicted in Fig 1.

**Fig 1.**
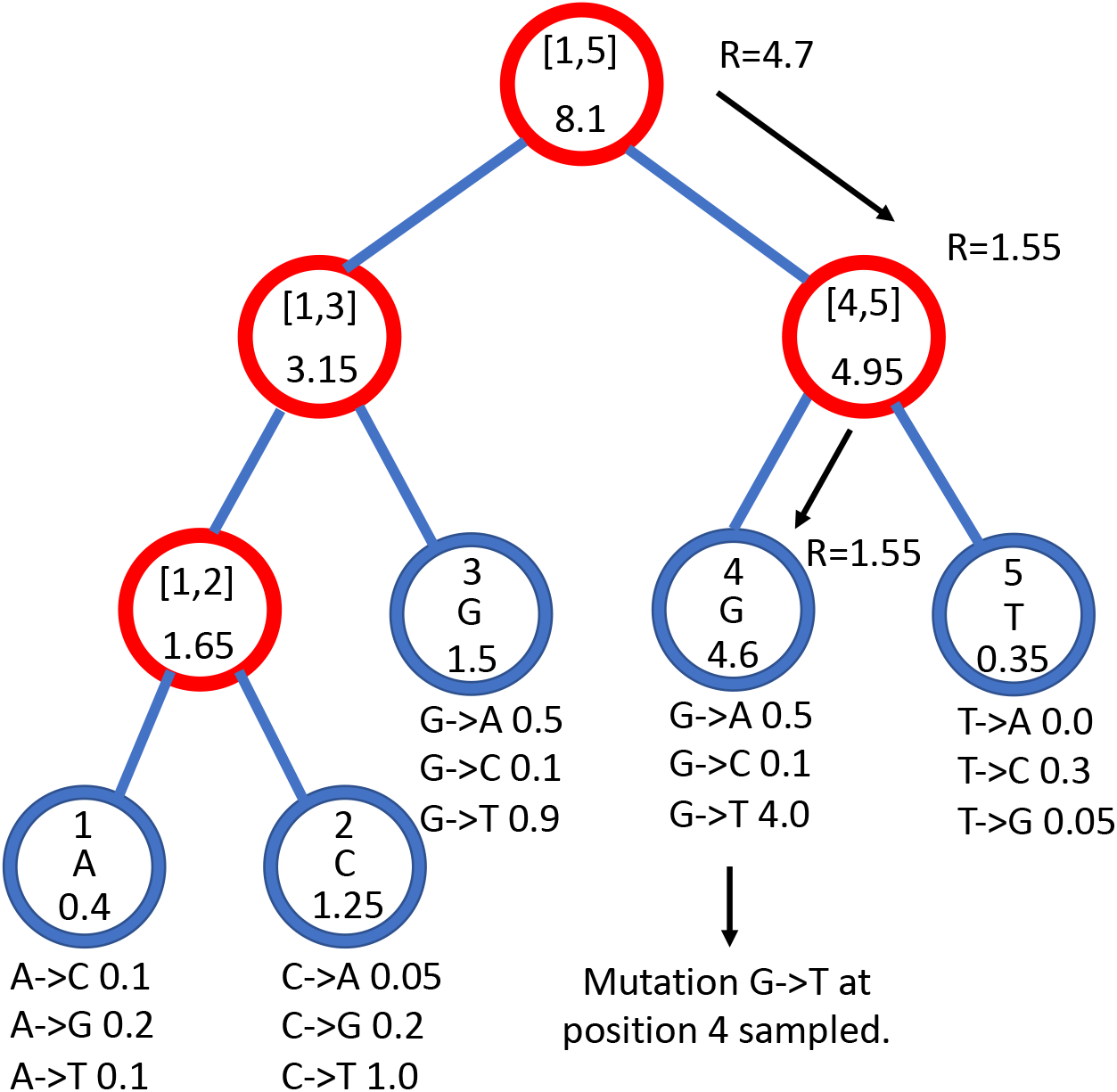
Example genome tree and genome tree search. An example genome search tree for ancestral genome ACGGT. Blue nodes are terminal and red nodes are internal. Inside each node we represent on top the genome positions represented by the node; at the center inside terminal nodes we show the allele of the node; at the bottom of nodes is their total rate. Under each terminal node we show the example relevant mutation rates. The black arrows show an example sampling of one mutation event. A parameter *R* is assigned an initial random number sampled uniformly between 0 and the total rate 8.1, in this case it is *R* = 4.7. As we move downward, the value of *R* can decrease, as described in Algorithm 2, determining which site will mutate and how. Here, an initial *R* = 4.7 results in the sampling of a G → T mutation at genome position 4.

Sampling a mutation time is done as in the vanilla approach: sampling from an exponential distribution with parameters determined by the total mutation rate at the root of the genome tree. Then, to sample a specific substitution event at a specific genome position, we first sample a random value uniformly in [0, 1) and multiply it by the total mutation rate *R*. Then, we traverse the tree from the root to the terminal node corresponding to the mutated position, which takes log(*L*) time. Finally, once reaching the corresponding terminal node (genome position) we choose a random substitution event affecting this position and correspondingly a new allele *a* for this position. An example mutation sampling is depicted in Fig 1. A pseudocode description of this algorithm is given in Algorithm 2. The cost of this approach is linear in the number of alleles, making it much more efficient than classical simulation methods based on matrix exponentiation when large state spaces (e.g. codon models) are considered. Furthermore, the computation cost for simulating under a codon model can be further reduced by considering that typically a codon model only allows a maximum of 9 substitution events from any codon, so at each terminal node we only need to consider a maximum of 9 events and rates at any time. Thanks to this, the cost of running a codon model with this approach is similar to the cost of running a nucleotide model.

##### Algorithm 2

Sampling of a substitution event along a genome tree, and updating the genome tree.

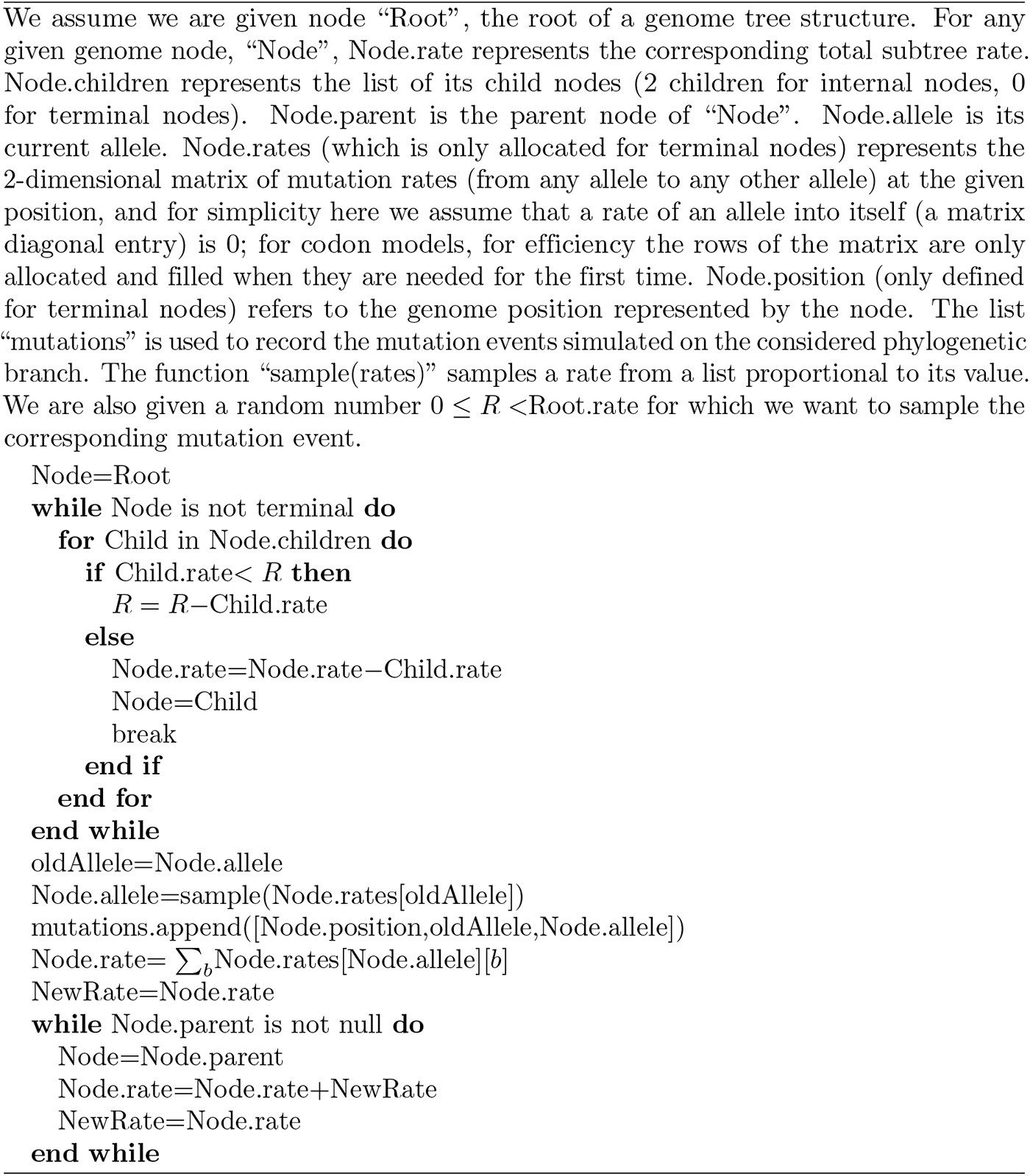

As mentioned before, once a mutation event is sampled, we need to modify the sampling process so that the change in allele at the mutated position is taken into account, since this change usually affects local and global mutation rates (a rare exception is for example when substitution rates are all equal). Modifying our genome tree following a substitution event is both simple and efficient: we simply need to modify the rates and allele at the mutated terminal node, and then update the rate of all ancestors of this terminal node accordingly. Algorithm 2 for example describes how to sample a substitution event from a genome tree as well as how to update the genome tree accordingly. Again, this can be done in log(*L*) time for each new substitution event sampled. However, while this is efficient for simulating evolution along a temporal line, that is, along a single branch of the phylogeny descendant from the root, it becomes inefficient for simulating evolution along a phylogenetic tree. This is because, if we modify the tree, then we cannot use it as it is for the sibling nodes. In other words, when we reach a split in the phylogenetic tree, and we have two children of the same phylogenetic node, we need to pass the same genome tree of the phylogenetic parent node to both phylogenetic children. However, we can’t only pass a pointer to the same tree to both children, because evolving along one branch leading to one sibling would modify the genome tree also for the other sibling. If we take the approach of duplicating the genome tree at each phylogenetic split, we end up with a cost *O*(*NL*), which we are trying to avoid. For this reason, we devise an alternative, hierarchical, multi-layered approach to evolving a genome tree, described below. Later on in the text we also describe the extension of our approach and of the genome tree structure to simulate indels.

#### Hierarchical, multi-layer evolving genome tree

In order to use our genome tree structure to sample mutations along a phylogenetic tree, we add a further “vertical” dimension to it. At each branch of the phylogenetic tree, instead of modifying a genome tree, we take the approach of building on it, without modifying the starting genome tree nodes, so that the original genome tree is not lost but instead is preserved at “layer 0” of our multi-layer structure. When we sample a mutation, we create a few new genome tree nodes in the corresponding layer of the structure, instead of not an entire new genome tree. By doing this, we can effectively adapt a (multi-layer) genome tree as new mutations are sampled without losing the original genome tree. This means that when we can pass the same genome tree to two children of a phylogenetic node without needing to duplicate the genome tree structure. Instead, we simply remove (de-allocate, or ignore) the genome tree nodes that have been added to other layers by the descendants of the first child node, and pass the same genome tree structure to the two considered child nodes. A graphical representation of an example multi-layer genome tree and its evolution is given in Fig 2.

**Fig 2.**
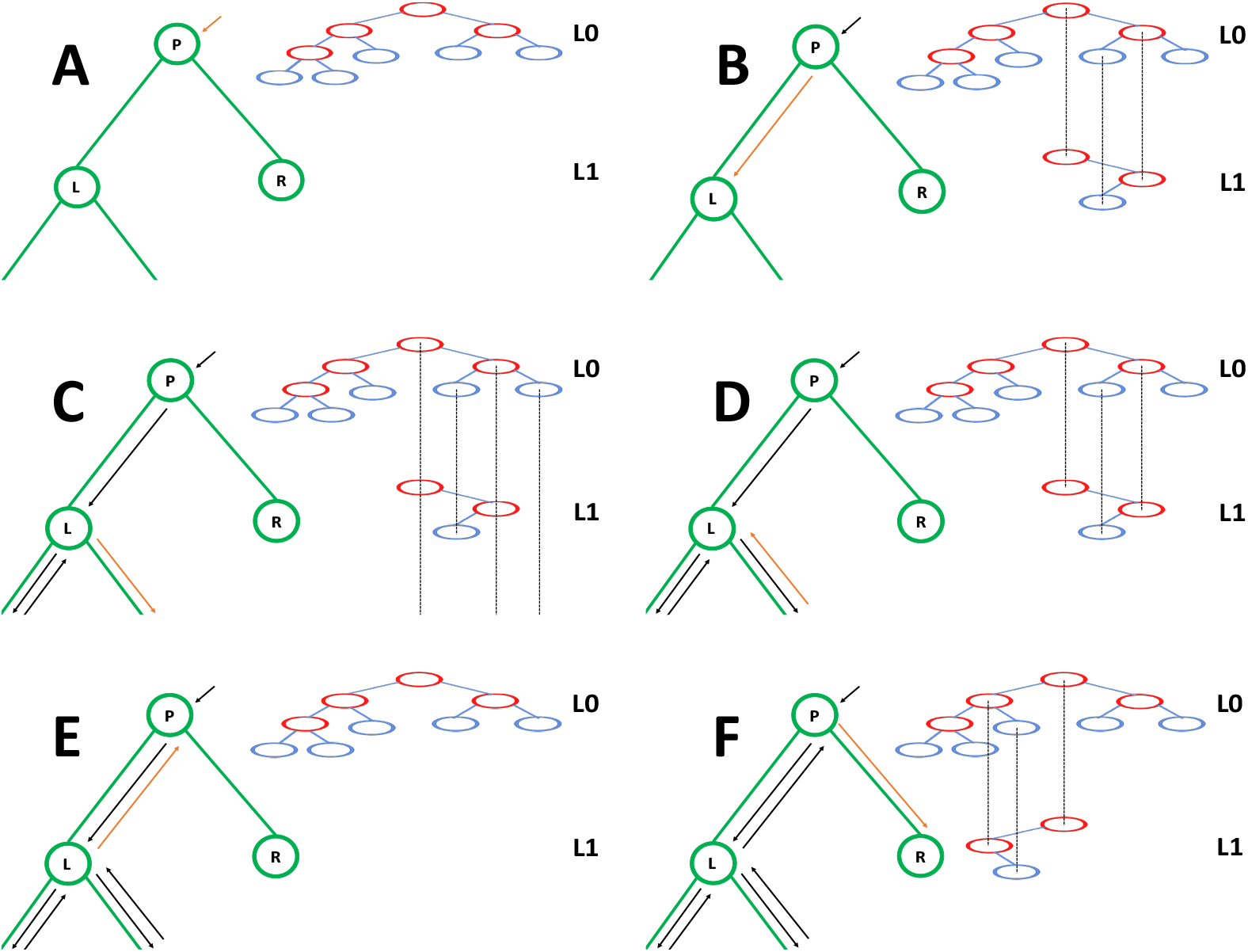
Example of multi-layer genome tree and its evolution. We track the evolution of the multi-layer genome tree starting from the genome tree of Fig 1. Colors for the genome tree are the same as in Fig 1. In green, on the left side of each panel, we show an extract of the phylogenetic tree containing three nodes (“P” for parent, which in this example is the root of the phylogeny, and “L” and “R” for left and right node). “L” has further descendants, but we don’t show them here and only focus on this triplet of nodes as an example. The orange arrow along the phylogenetic tree shows the current step of the preorder traversal being considered by the given panel. Black arrows show past steps. Vertical dashed black lines in the multi-layer genome tree connect nodes that represent the same portions of the genome but that are in different layers. “L0” stands for “Layer 0” and “L1” for “Layer 1”. **A** At the phylogenetic root “P” we initialize the genome tree for layer 0. **B** As we move to child “L”, a new substitution is sampled (as in Fig 1) and 3 corresponding genome nodes are created in layer 1. These nodes correspond to the nodes in the original genome tree whose rate is affected by the new mutation. **C** As we traverse the subtree of the descendants of L, new nodes and mutations might be added in the layers below. **D** We are finished traversing the subtree of the descendants of L, and we return to L, at which point all nodes in layer below 1 have either been removed or have become irrelevant. **E** We return to P, at which point the genome tree nodes previously added layer 1 are also ignored or deleted. **F** We move from P to R, and in doing so new mutation events might be sampled and the corresponding genome nodes might be added to layer 1 (new genome tree nodes corresponding to 1 new substitution are shown in the new layer 1).

We start with a genome tree at the phylogenetic root node; additional nodes are then added at further layers. A genome tree layer *n* represents the genome nodes specific to a particular depth of the phylogenetic tree; phylogenetic nodes closer to the root (in terms of number of branches that need to be traversed from the root) will be associated with a lower *n*, and those more distant from the root with higher *n*. All the initial nodes of the original genome tree belong to layer 0, the layer corresponding to the phylogenetic root. Then, as we move from the phylogenetic root to its first child, we add nodes to the tree in layer 1, representing the consequences of mutation events happening along the branch between the phylogenetic root and the first child. Nodes in layer 0 only point to nodes in layer 0, and never to nodes in other layers. More generally, nodes in layer *n* only point to nodes in layers *m* ≤ *n*. Every time the multi-layer genome tree is passed from phylogenetic parent (layer *n*) to child (layer *n* + 1), new nodes are added to the corresponding layer (*n* + 1) if mutation events occur on the corresponding phylogenetic branch.

We traverse the phylogenetic tree in preorder traversal, so, starting from the root, we move to the first child, to which we pass the initial genome tree, add new layers, then do the same for this child’s children. For each new mutation occurring on this phylogenetic branch connecting the root and its first child, we traverse the genome tree, and every time we would modify the genome tree (to update the mutation rates following a change of allele at a position) we instead create new genome tree nodes in the child layer. Once we have traversed the whole phylogenetic subtree of the first child of the root, we have to move to second child of the root. This operation does not incur the cost of duplicating any part of the genome tree, as we only need to pass to the second child the pointer to the root of layer 0 of the hierarchical genome tree. Similarly, at any internal phylogenetic node at layer *n*, to both children we pass the pointer to the root of layer *n* of the genome tree. The only additional step which might be required is the de-allocation of nodes in layer *n* + 1 as we move from one node to its sibling (thanks to our preorder traversal, the nodes currently in and below this layer will not be used again), but this step at most only slows simulations by a small constant factor.

At the start of the simulations for each branch, moving from layer *n* to *n* + 1, we first create a new genome root node for layer *n* + 1. This root initially points to the same children as the genome root at layer *n*, and it also has the same total rate. After creating a new layer root, we sample mutation events for the current phylogenetic branch. To sample mutations, we follow the binary search tree determined by the root of layer *n* + 1. As a new mutation event is picked, we either create new layer *n* + 1 nodes, or modify existing layer *n* + 1 nodes. When sampling a new mutation, every time we reach a node in the genome tree, we either modify the rate of the node, if it’s in layer *n* + 1, or we create a new layer *n* + 1 node, if the original node was in a different layer. The new node is given at first the same children as the original node. When a terminal node is reached, we calculate its new rates (unless they have already been created before for some other node in the phylogenetic tree, in which case we just retrieve them from a dictionary) and total rate, and we pass the new total rate to its parent node, which uses it to update its own total rate, and so on. In total, the cost of sampling a new mutation event and updating the multi-layered structure is *O*(log(*L*)). A sketch of the mutation sampling process and multi-layer genome tree update is given in Algorithm 3. The total cost of the algorithm is then *O*(*L* + *N* + *M* log(*L*)), where the addendum *L* is due to the initial creation of the layer 0 genome tree, and *N* is due to the tree traversing process. In a scenario like SARS-CoV-2 genomic epidemiology, this can lead to dramatic reduction in computational demand compared to the standard *O*(*NL*) in the field, since *M* appears typically not distant from *N* [22–24]. We give a summary of the global hierarchical method in Algorithm 4.

##### Algorithm 3

SampleMutation(Node,Layer,*R*): Sampling of a mutation event along a multi-layer genome tree.

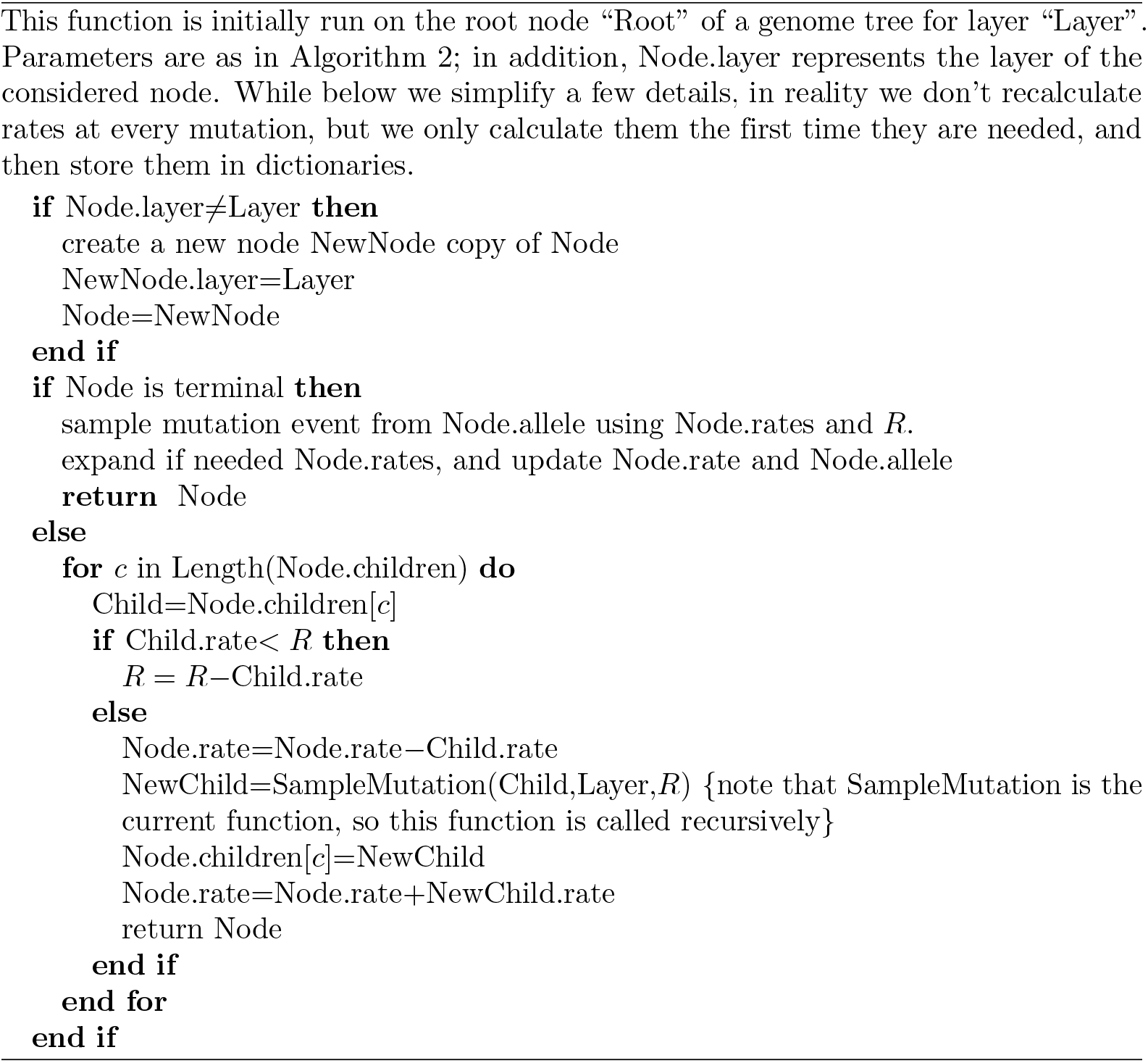

##### Algorithm 4

SimulatePhyloNode(Node,GenomeNode,Layer): Hierarchical algorithm for simulating sequence evolution along a branch of the phylogenetic tree.

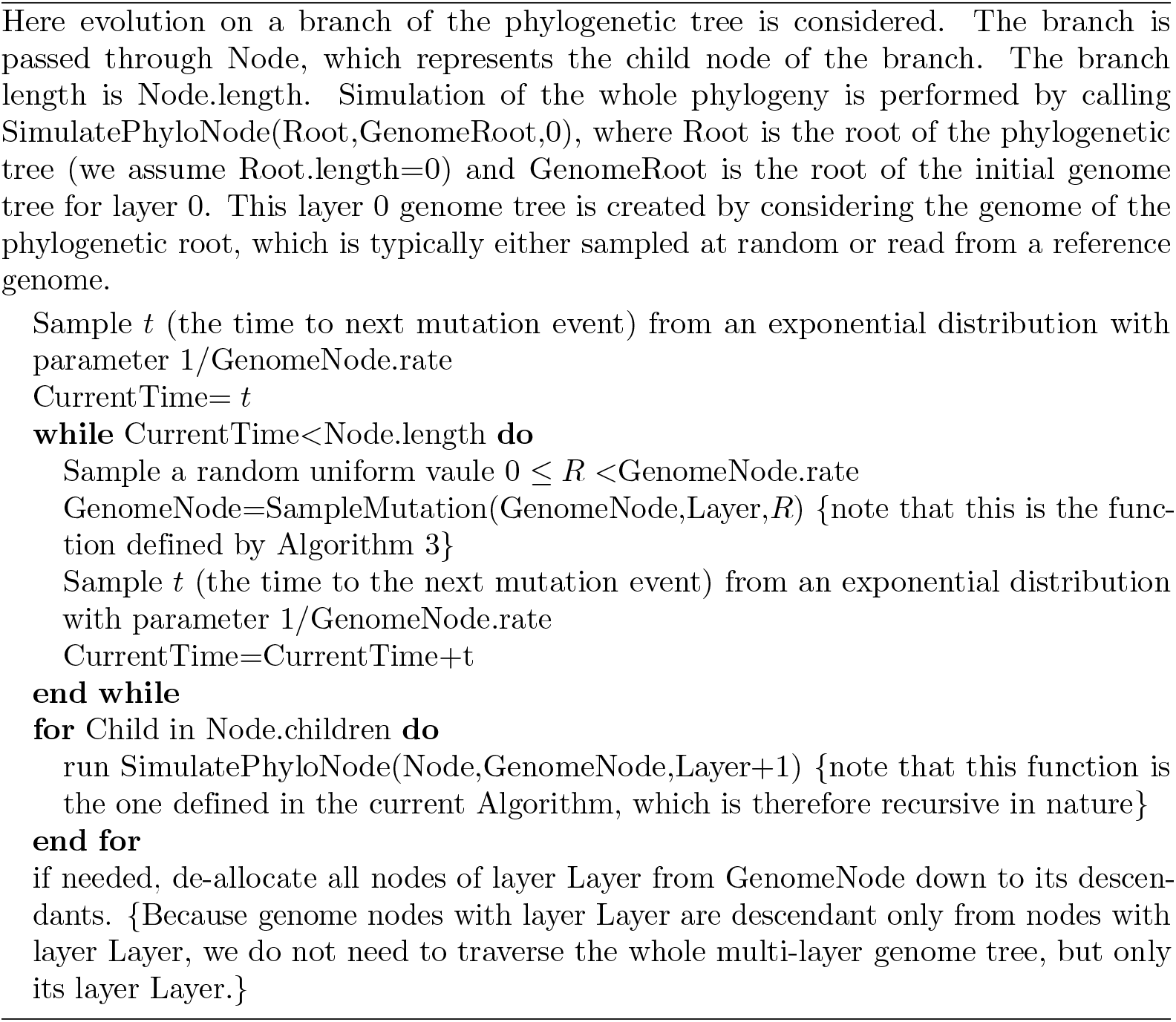

### Indels

We further extended the multi-layer genome tree approach to efficiently simulate insertions and deletions. Each leaf on the genome tree is assigned a deletion rate, insertion rate, and substitution rate, denoted *R*_*d*_, *R*_*i*_, and *R*_*s*_ respectively, and the total mutation rate for the leaf will be *R*_*d*_ + *R*_*i*_ + *R*_*s*_. The substitution rate *R*_*s*_ itself is the sum of all substitution rates from the current allele of the leaf. Insertions are modeled as occurring on the right (3′ end) of the sampled position; to model insertion at the 5′ end of the genome, a dummy terminal genome tree node is employed representing the leftmost end of the genome, and is initialized with *R*_*s*_ = *R*_*d*_ = 0 but with non-zero *R*_*i*_. Just as with the substitution rates, which can be site specific, *R*_*d*_ and *R*_*i*_ can be drawn from a gamma distribution, or can be constant across the genome. When a mutation event is sampled at a node, it will be sampled as a deletion, an insertion, or a substitution proportionally to *R*_*d*_, *R*_*i*_ and *R*_*s*_.

Our software allows for indels with lengths drawn from a number of parametric distributions following the options allowed with INDELible, see Table 1 for an overview of the various distributions that have been implemented. Sampled indels have always length ≥ 1.

**Table 1.** Indel length distribution options.

| Distribution | Parameters | $\mathbf{P}(X = n), n > 0$ |
| --- | --- | --- |
| Geometric | $p$ | $(1 - p)^{n-1}p$ |
| Negative Binomial | $p, k$ | $\binom{k+n-1}{n}(1 - p)^{n-1}p^k$ |
| Zeta | $a$ | $n^{-a}/\zeta(a)$ |
| Lavalette | $a, k$ | $(\frac{kn}{k-n+1})^{-a}$ for $n \leq k$ |
| Discrete | vector $\mathbf{v}$ | $v_n$ |

Below we explain in more detail the algorithm used to efficiently simulate insertions and deletions using multi-layered genome trees. In short, insertion events are simulated by adding new small subtrees to the genome tree in the current layer. Deletion events are instead simulated by setting substitution and indel rates to 0 in deleted nodes.

#### Insertion Algorithm

The algorithms for simulating insertions and deletions mostly proceed as the one simulating substitutions (Algorithm 3) in that we traverse the genome tree to find the terminal node “Leaf” affected by the next sampled mutation. We then sample the type of the next mutation event (insertion, deletion, or substitution) proportional to the corresponding mutation rates *R*_*i*_, *R*_*d*_ and *R*_*s*_ of “Leaf”. The process for simulating a substitution remains the same as before. If instead a new insertion event is simulated, we sample a length *l* for the inserted material from the corresponding prior distribution, and then add a new subtree to the genome tree as detailed in Algorithms 5 and 6.

##### Algorithm 5

insertNode(Node, *l*): this function inserts a new genome subtree at the given terminal genome tree node “Node” at which the insertion is sampled, given the insertion length *l*. We assume that Node is part of the current layer, and that new nodes are created at the current layer.

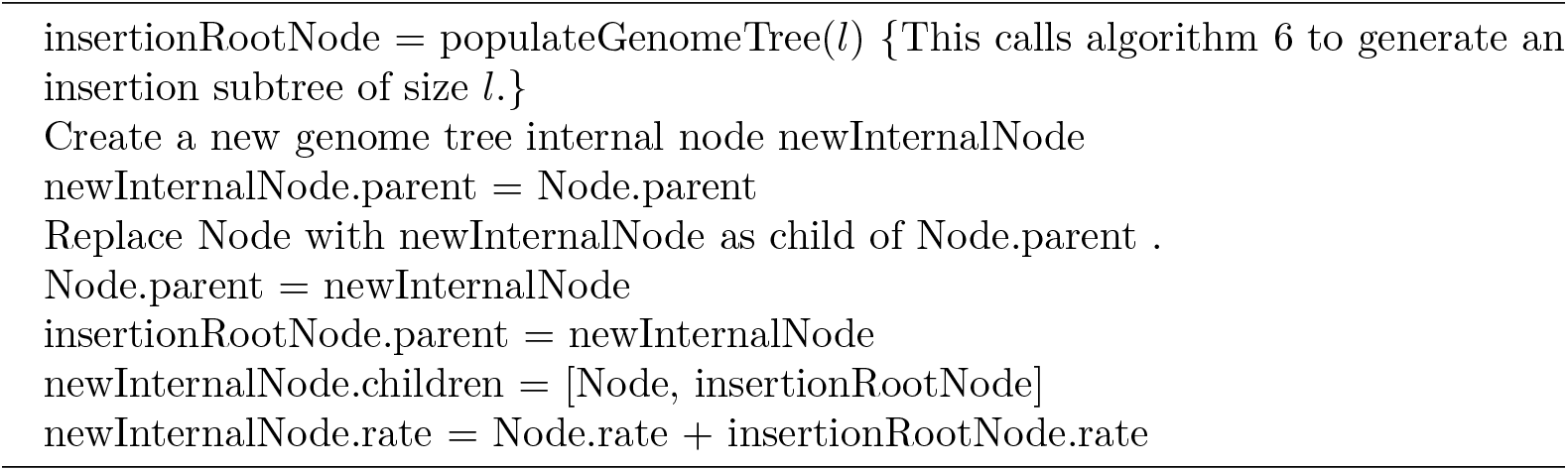

Note that the addition of new subtrees to the genome tree will typically make it less balanced, and a potentially less efficient search tree. In typical scenarios considered here, that is, when divergence is low and all genomes are closely related to the root, the effect of this imbalance on the overall search-efficiency of the genome tree will be extremely minor.

#### Deletion Algorithm

If the next mutation event at Leaf will instead be a deletion, again, we first sample a deletion length *l*, and then we proceed to set to 0 the total mutation rate for node Leaf and its following *l* − 1 positions of the genome in the current layer, ignoring positions

#### Algorithm 6

populateGenomeTree(*l*): this function recursively creates a new genome subtree for a given insertion of length *l*, and returns the root node of the subtree.

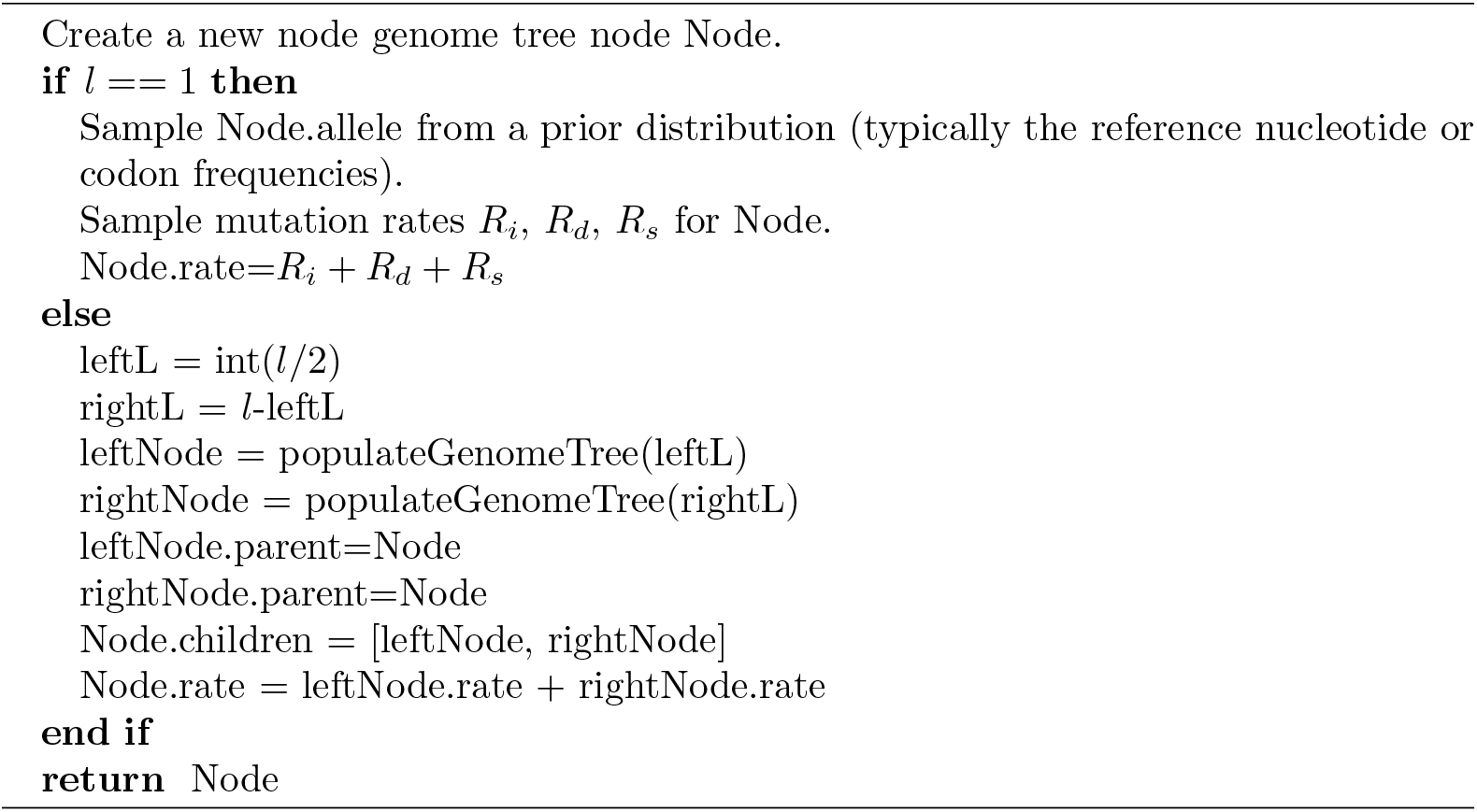

that are already deleted. The main subtlety of this approach is to avoid traversing the whole genome tree (incurring a cost of *O*(*L*) operations), to delete these *l* characters. Algorithm 7 below performs this task efficiently employing at most *O*(*l* log(*L*)) operations. This algorithm returns the number of characters deleted. If Leaf represents a position near the 3′ end of the genome, the sampled deletion length might overflow past the end of the genome, and so fewer than *l* positions might be effectively deleted.

##### Algorithm 7

deleteNodes(Rand,GenomeNode,Layer,RemainingDeletions): recursive algorithm for deleting nodes of the genome tree following a deletion event. It returns the number of positions deleted. “Rand” is the random number that has been used to sample the deletion event - here it’s used to direct the search to the first deleted positions and the following ones.

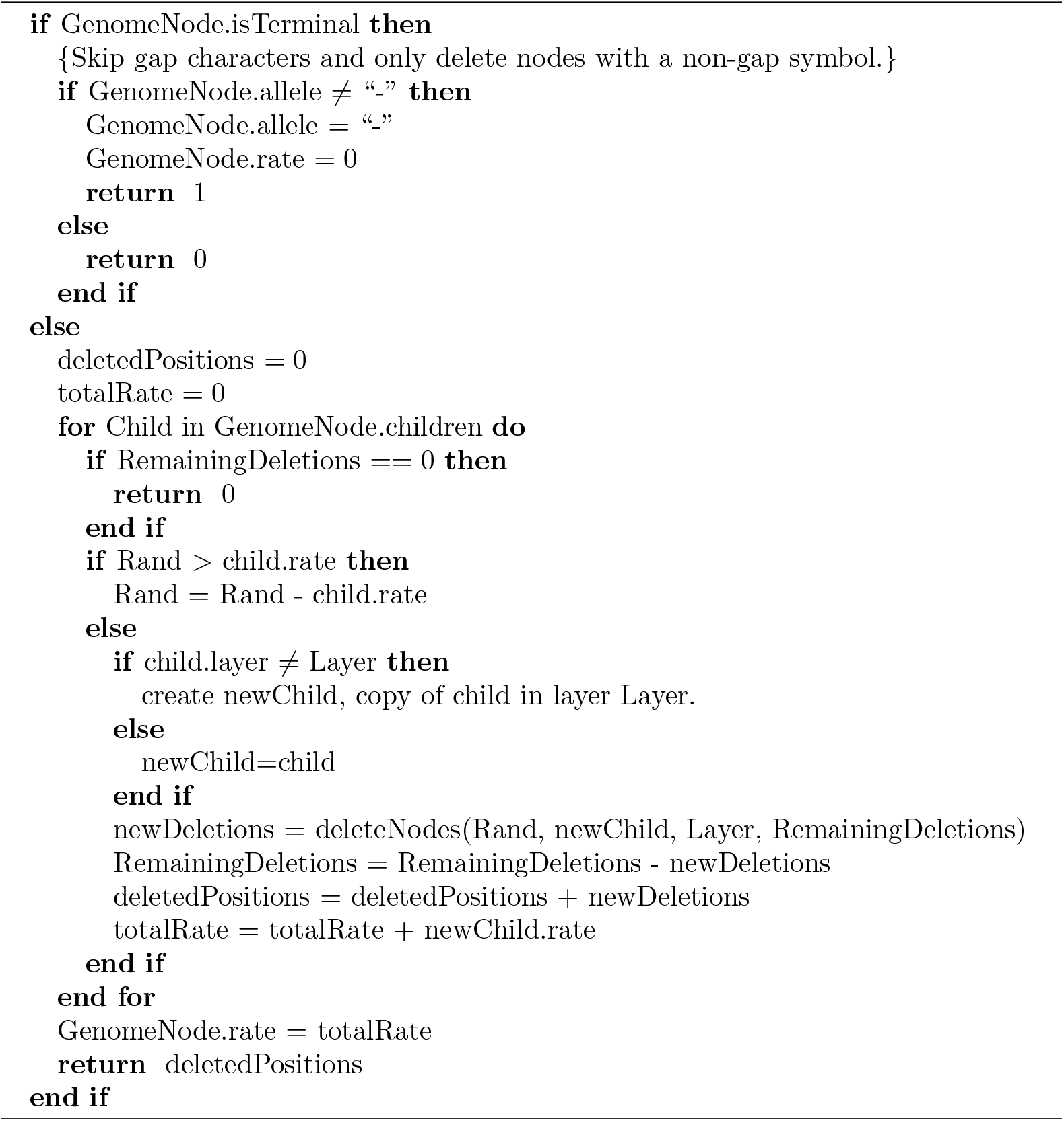

### Further details of the implementation

#### Substitution models

Thanks to our algorithm, we can allow any substitution model without incurring a dramatic increase in computational demand, and without risking numerical instability (which can sometimes be a problem with classical matrix exponentiation approaches). Users can easily specify different nucleotide substitution matrices (e.g. JC [28], HKY [34], or GTR [6]). By default, we adopt the most general nucleotide substitution model, UNREST [31], using as default rates those we estimated from SARS-CoV-2 [22].

We also implemented codon models, which, with our hierarchical approach, come at only a small additional computational demand compared to nucleotide models. To define substitution rates of codon models, we use an extension of the GY94 [35] model, and separately model the nucleotide mutation process and the amino acid selection one. Unlike GY94 (which assumes an HKY nucleotide mutation process), we allow any general nucleotide mutation process as defined by an UNREST matrix. Then, nonsynonymous mutations rates are modified by a single factor *ω* (see next section for variation of *ω* across codons). Under this model, a substitution from codon *c*_1_ to codon *c*_2_ therefore has rate:

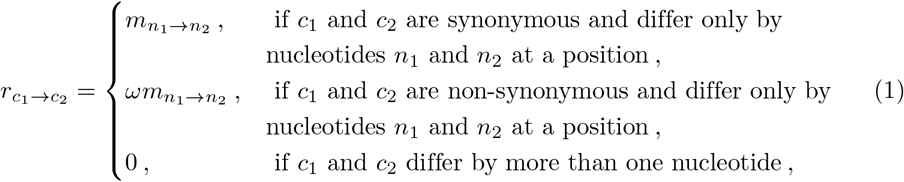

where 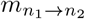 is the mutation rate from nucleotide *n*_1_ to nucleotide *n*_2_.

We don’t allow, at this stage, instantaneous multi-nucleotide mutation events, or amino acid substitution models, but we plan to address them in future extensions. A description of currently implemented models and a comparison with those in other similar simulation software is given in Table 2.

**Table 2.**
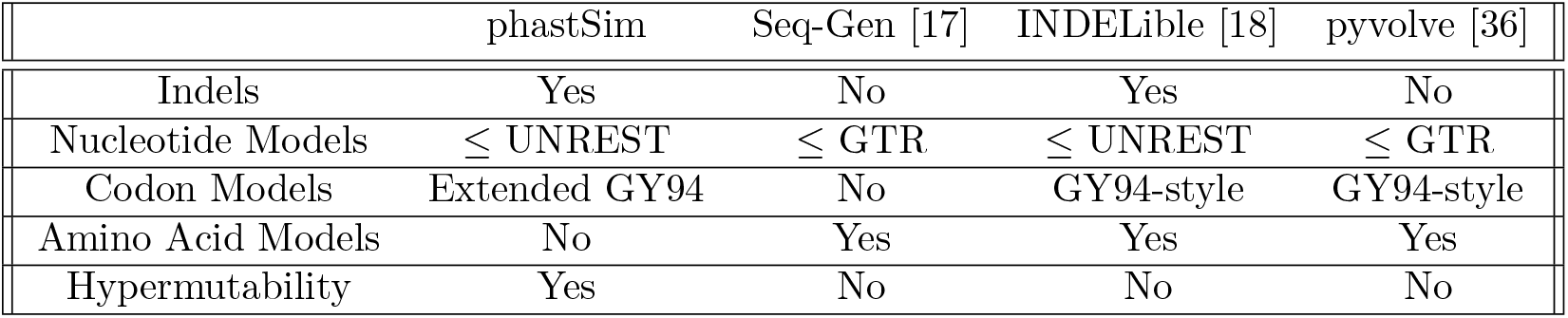
A comparison of features of different sequence evolution simulation software packages.

### Models of rate variation

We consider four types of variation in rates across the genome. These types can be used in combination, or separately, as required.

The first type of variation is changes in the position-specific mutation rate across the genome. Every nucleotide position *i* in the genome (even when using a codon model) is assigned its own mutation rate scaling factor *γ*_*i*_. This means that, at position *i*, the mutation rate from any nucleotide *n*_1_ to any other nucleotide *n*_2_ becomes 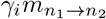. We allow two ways to sample values of *γ*_*i*_ for each *i*. One way is to sample them from a continuous Gamma distribution with parameters Γ(*α, α*), with *α* specified by the user; this results in each genome position having a distinct *γ*_*i*_. Alternatively, we allow the definition of discrete categories, with a finite number of categories, each with its own proportion of sites and *γ* rate.

The second type of variation we model is variation in *ω*, with each codon position *i* across the genome being given its own *ω*_*i*_. As with *γ*_*i*_, values of *ω*_*i*_ can either be sampled from a continuous Gamma or a finite categorical distribution.

Lastly, to accommodate the strong variation in mutation rates observed in SARS-CoV-2 [22, 23] attributable to APOBEC, ADAR, or ROS activity, we introduce a new model of rate variation. This model allows, for a certain position, to have one specific mutation rate (from one specific nucleotide to another specific nucleotide) enhanced by a certain amount *μ*. In this case we only allow a categorical distribution, with the first category having no enhancement (*μ* = 1) and the other categories having *μ* < 1. For any nucleotide position *i* that is assigned a hypermutable category and therefore has *μ*_*i*_ > 1, we then sample uniformly a start nucleotide *n*_*s*_ and a destination nucleotide *n*_*d*_. The mutation rates 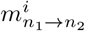 for position *i* then become:

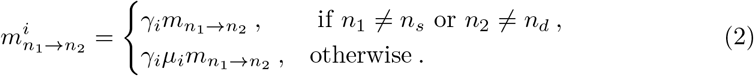

#### Rate normalization

We assume that, for the given input phylogenetic tree, branch lengths represent expected numbers of substitutions per nucleotide - no matter if a nucleotide or a codon model is used. However, as mutations accumulate across the phylogeny, the total mutation rate of the genome might slightly change. This is because we allow substitution models that are not reversible and not at equilibrium. Therefore, we always consider the mutation rates at the root genome to normalize the mutation rates. This means that while branch lengths near the root accurately represent the expected numbers of substitutions per nucleotide, lower down the tree this might slightly change.

#### Output formats

As default, our software creates an output file where it stores information about which genome position evolved under which rate. It also creates a file where each tip name is listed together with the mutations it contains that distinguish its genome from the reference genome. In scenarios similar to SARS-CoV-2 datasets (where each genome is very similar to the reference), this format requires much less space and time to generate than FASTA or PHYLIP formats (see the “vanilla” approach subsection).

An optional output format that our software can create is a tree in Newick format, where each branch of the input phylogeny is annotated with a list of mutation events that occurred on that branch. This format is richer than the others, as it provides information regarding each mutation event, even those that might be over-written by other mutations at the same position; it is also more efficient than multiple sequence alignment formats in the scenario of short branch lengths considered here. We also allow a binary analogue of this annotated Newick tree, called a MAT (mutation annotated tree) [37], which is compatible with the phylogenetic software UShER [27].

Finally, we also allow the creation of unaligned FASTA output. However, note that the creation of a FASTA file costs *O*(*NL*) in time and space. In the case simulations are performed without indels, we also allow the generation of a PHYLIP format alignment output.

#### Python package

Our software phastSim is implemented as a Python package, and can be found at https://github.com/NicolaDM/phastSim or https://pypi.org/project/phastSim/. phastSim uses the ETE3 library [38] to robustly read input trees in different variants of the Newick format.

## Results

### SARS-CoV-2 scenario

To assess the performance of our approach compared to existing evolutionary simulators, we consider different scenarios typical for genomic epidemiology. First, we consider the simulation of a scenario similar to SARS-CoV-2 evolution. We simulate trees with a custom script and under a Yule process with birth rate equal to genome length (29,903), so to have in the order of one mutation per branch. Evolution is simulated under an UNREST model [31] with rates inferred from SARS-CoV-2 data [22] where possible (for phastSim, pyvolve [36] and INDELible [18]) and a GTR model [6] otherwise (for Seq-Gen [17]). We run INDELible both using method 1 (“INDELible-m1”, which uses matrix exponentiation to model substitutions) and method 2 (“INDELible-m2”, which instead uses the Gillespie approach for the same task). For now we ignore sequence rate variation. While phastSim and pyvolve are both Python implementations, therefore sharing similar benefits (high compatibility with other packages and ease of extensions) and draw-backs (reduced efficiency compared to some other languages), we see that the two approaches have dramatically different computational demands (Fig 3): simulating 50 sequences under pyvolve requires on average more time than simulating 500,000 in phastSim. We can also see that INDELible-m2 is marginally more efficient than INDELible-m1 in this scenario, due to the low number of mutations per branch. However, while phastSim and INDELible-m2 are both similar Gillespie approaches, simulating 5,000 sequences with INDELible-m2 requires slightly more time than simulating 500,000 sequences in phastSim (Fig 3), despite the fact that INDELible is coded in C++. Seq-Gen appears to be very efficient, but it’s still more than one order of magnitude slower than phastSim on large phylogenetic trees in this scenario. Also note that, for large trees considered here, we can reduce computational demand in phastSim by more than 5-fold by not producing a FASTA output alignment; this way we can also save very significant amounts of memory demand. Regarding small trees (< 10^4^ tips) most of the demand in phastSim is associated with initializing the simulations (loading packages and initializing the genome tree structure); these initialization costs do not depend on tree size, and instead depend on genome size, and they are why phastSim is relatively less efficient on small trees. If simulation on small trees are indeed of interest, these initialization costs could be reduced by re-using the same genome tree structure over multiple replicates, or, in the case of simple evolutionary models, by using our “vanilla” simulation approach.

**Fig 3.**
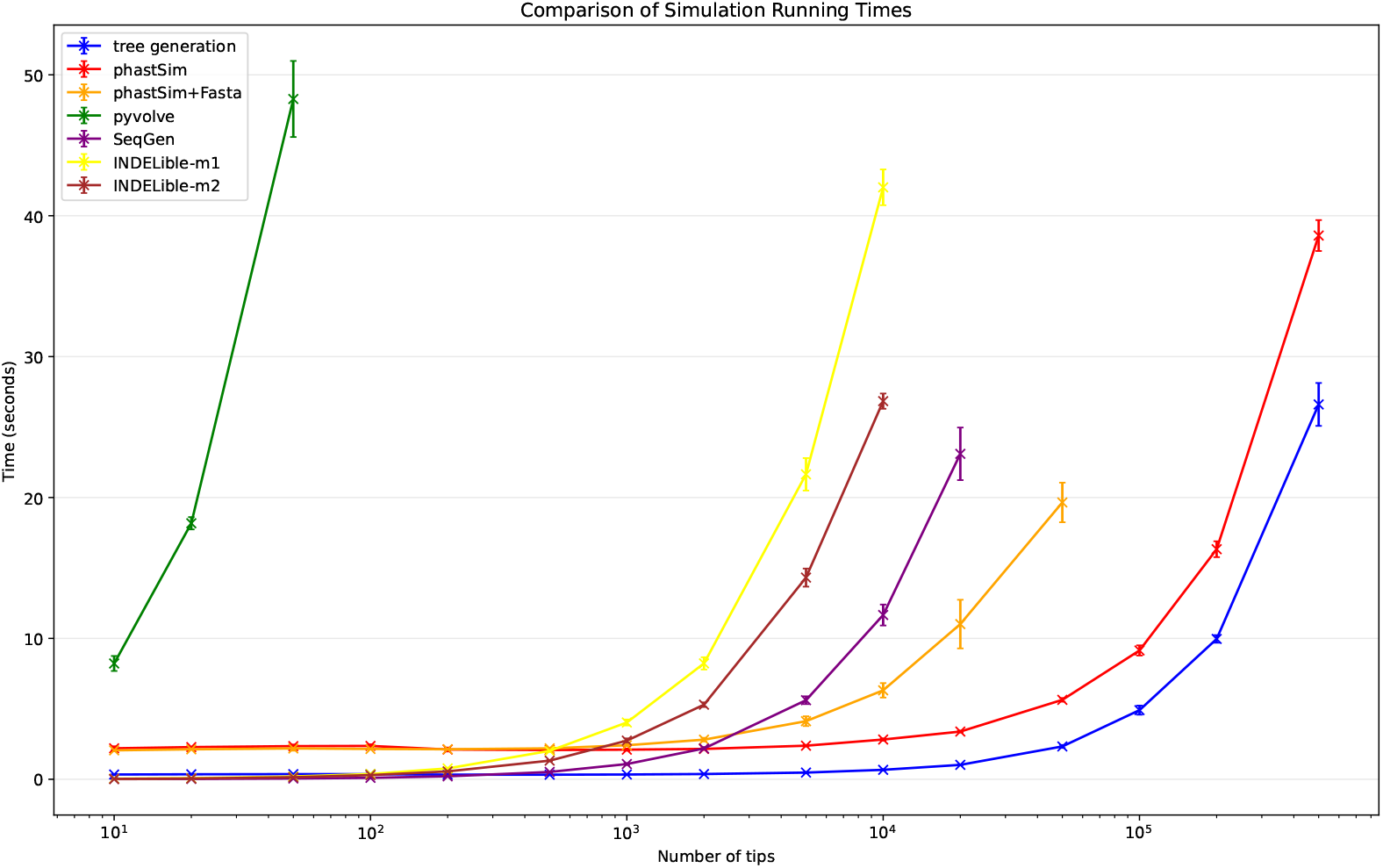
Comparison of running times of different simulators in a scenario similar to SARS-CoV-2 data. On the Y axis we show the number of seconds it takes to perform simulations using different software. On the X axis is the number of tips simulated. Each boxplot represents ten replicates. We do not run the most demanding simulators when each replicate would take substantially more than 1 minute to run. In blue is the computational demand for generating the random trees with a customised version of NGESH [39] distributed within the phastSim package; sequence simulation is performed conditional on these simulated trees. In red is the time to run phastSim with a concise output, and in orange is the time for phastSim with additionally generating a FASTA format output. In green is the demand of pyvolve, and in purple of Seq-Gen. In yellow and brown are respectively the time for running INDELible with method 1 (matrix exponentiation) and method 2 (Gillespie approach).

### Bacterial scenario

To demonstrate a scenario in which we are interested in simulating bacterial genome evolution within one outbreak, we use the *E. Coli* reference genome (https://www.ncbi.nlm.nih.gov/nuccore/U00096.3 [40], 4,641,652 nucleotides) as our root genome sequence. For now we still focus on the simple scenario of a nucleotide model without rate variation. We again assume a scenario typical for genomic epidemiology, that the birth rate of the simulated tree is equal to the genome length. The number of mutations simulated is therefore comparable to the number of branches in the tree.

As genome size increases, time and memory demand of traditional simulators is expected to grow linearly. Indeed, we now see that Seq-Gen takes considerably more time to simulate the same number of genomes than in the SARS-CoV-2 scenario (Fig 4). phastSim also has an increased computational demand, but only in terms of the initial step of generating an initial genome tree. This initial cost is linear with respect to genome length, but does not increase with the number of samples or with the number of mutations simulated. In total, in this scenario phastSim can simulate sequence evolution along trees with more than 1000 times more samples than Seq-Gen. A further reduction in computational demand, in particular in terms of the initial cost of generating a genome tree, can be obtained by using the “vanilla” algorithm (Fig 4), which however comes at the cost of narrowing the choice of evolutionary models to less complex ones.

**Fig 4.**
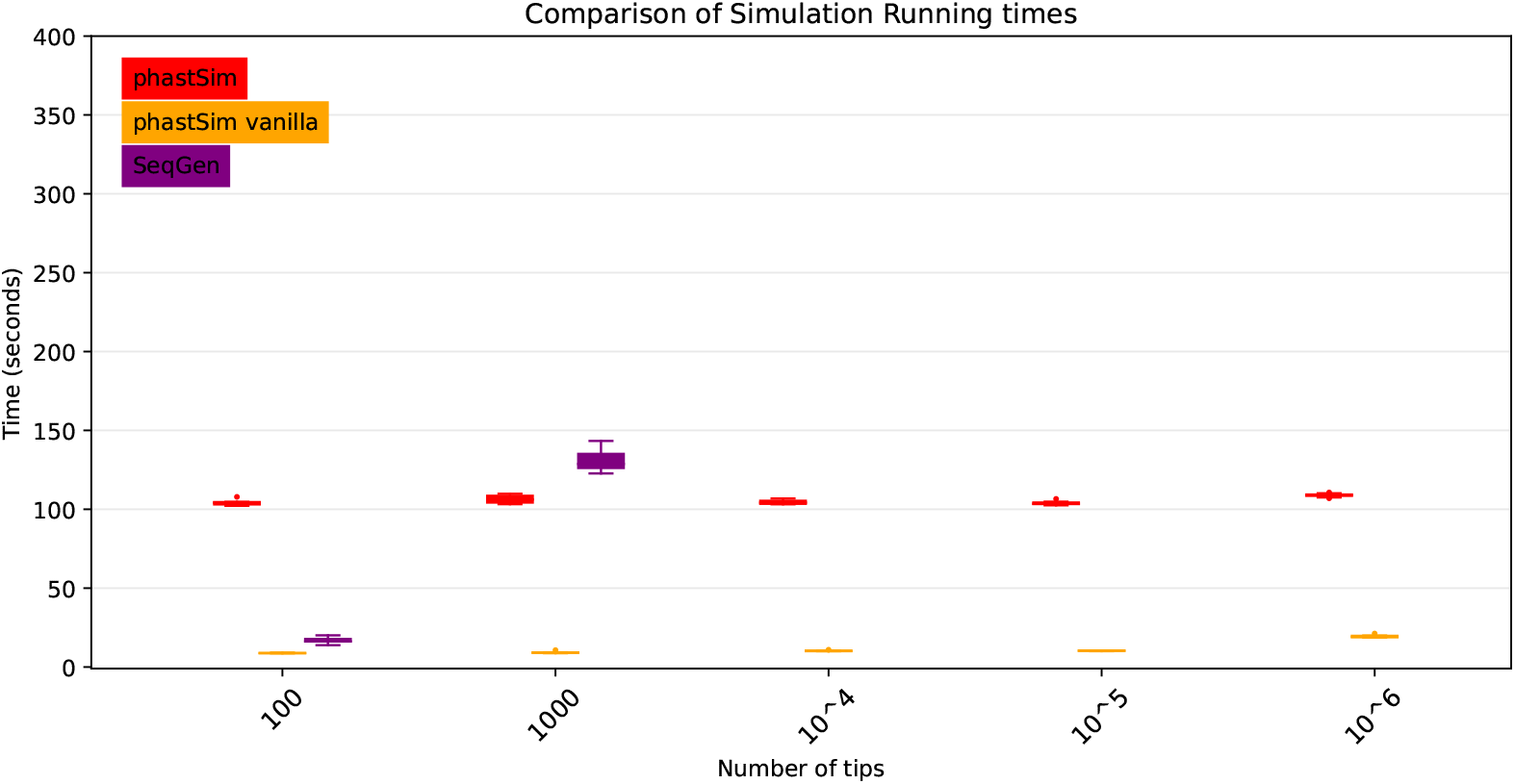
Comparison of running times of different simulators in a scenario similar to *E. Coli* outbreak data. On the Y axis we show the number of seconds it takes to perform simulations using different software. On the X axis is the number of tips simulated. Each boxplot represents ten replicates. We do not run Seq-Gen for more than 1000 tips due to high computational demand. In red is the time to run phastSim, and in orange is the time for phastSim with the “vanilla” approach. In purple is the time demand of Seq-Gen.

### Evolutionary and Indel models

One of the advantages of the approach we present here is that simulating evolution under increasingly complex models comes at almost no additional computational cost (Fig 5). It can be seen, for example, that INDELible-m1 and Seq-Gen incur a significantly higher cost when using a continuous variation in mutation rate, and that INDELible-m2 is more demanding when simulating discrete rate categories. Surprisingly, running INDELible with a codon model appears to come with no additional computational demand, similarly to phastSim (Fig 5). For these comparisons we have considered the SARS-CoV-2 simulation scenario.

**Fig 5.**
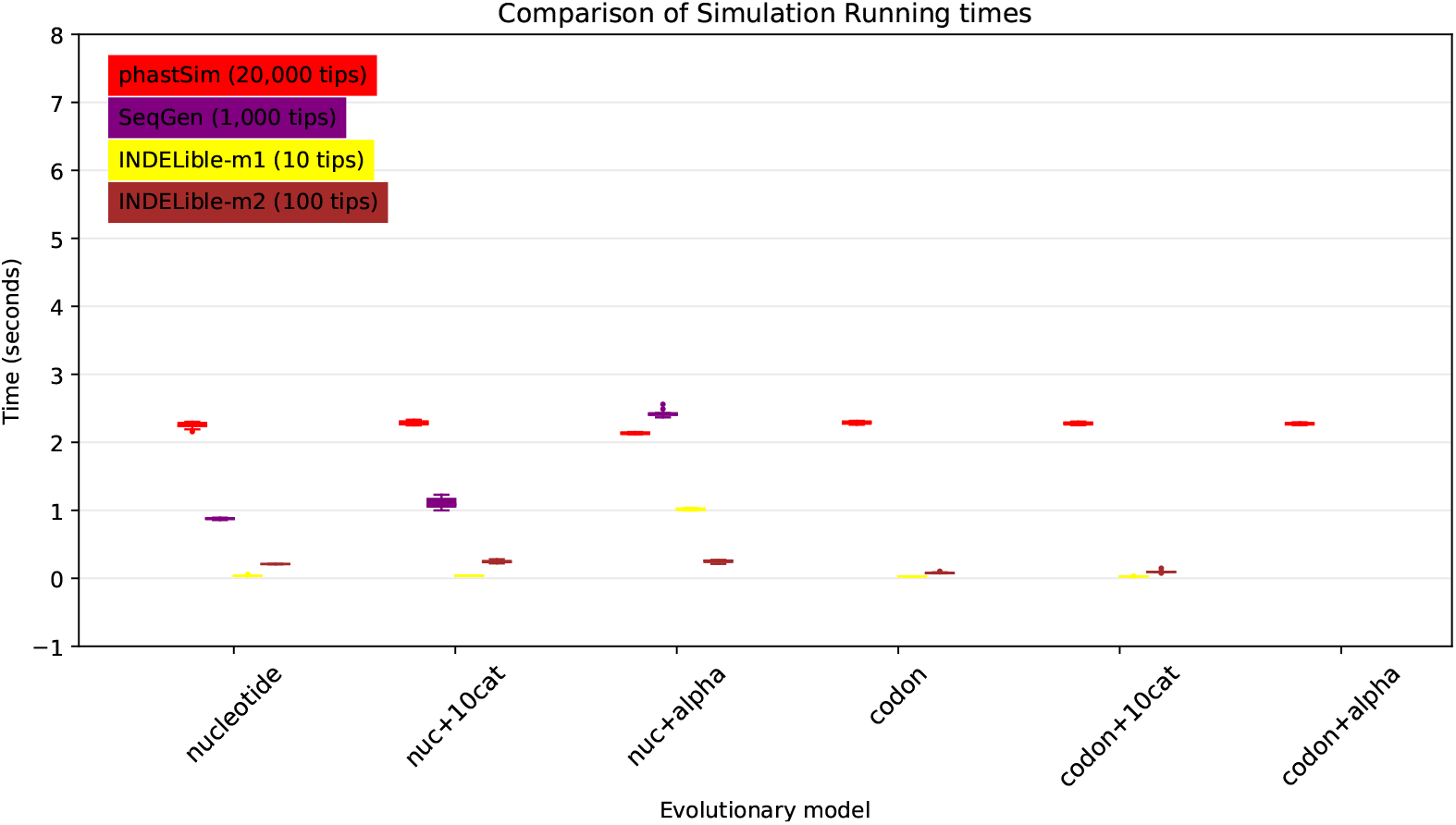
Comparison of running times of different simulators in a SARS-CoV-2 scenario using different evolutionary models. On the Y axis we show the number of seconds it takes to perform simulations using different software. On the X axis is the model used for simulations: “nucleotide” is a nucleotide substitution model without variation; “nuc+10cat” is a nucleotide model with 10 rate categories; “nuc+alpha” is a nucleotide model with continuous variation in rate (each site has a distinct rate sampled from a Gamma distribution); “codon” represents a codon substitution model; “codon+10cat” represents a codon substitution model with 10 categories for *ω*; “codon+alpha” is a codon model with continuous rate variation in mutation rate and in *ω* (only allowed in phastSim). Each boxplot represents ten replicates. Seq-Gen does not allow codon models. Colors are as in Fig 3. To aid the visual comparison, we use trees of different sizes for different simulators: 10 tips for INDELible-m1; 100 tips for INDELible-m2; 1,000 tips for SEq-Gen; 20,000 tips for phastSim.

Our algorithm also allows efficient simulation of insertion and deletion events (indels). Among the other simulators considered here, only INDELible can simulate indels. In the SARS-CoV-2 scenario, phastSim can simulate substitutions and indels for considerably larger phylogenies (about 10 times larger) than INDELible for the same computational run time (Fig 6).

**Fig 6.**
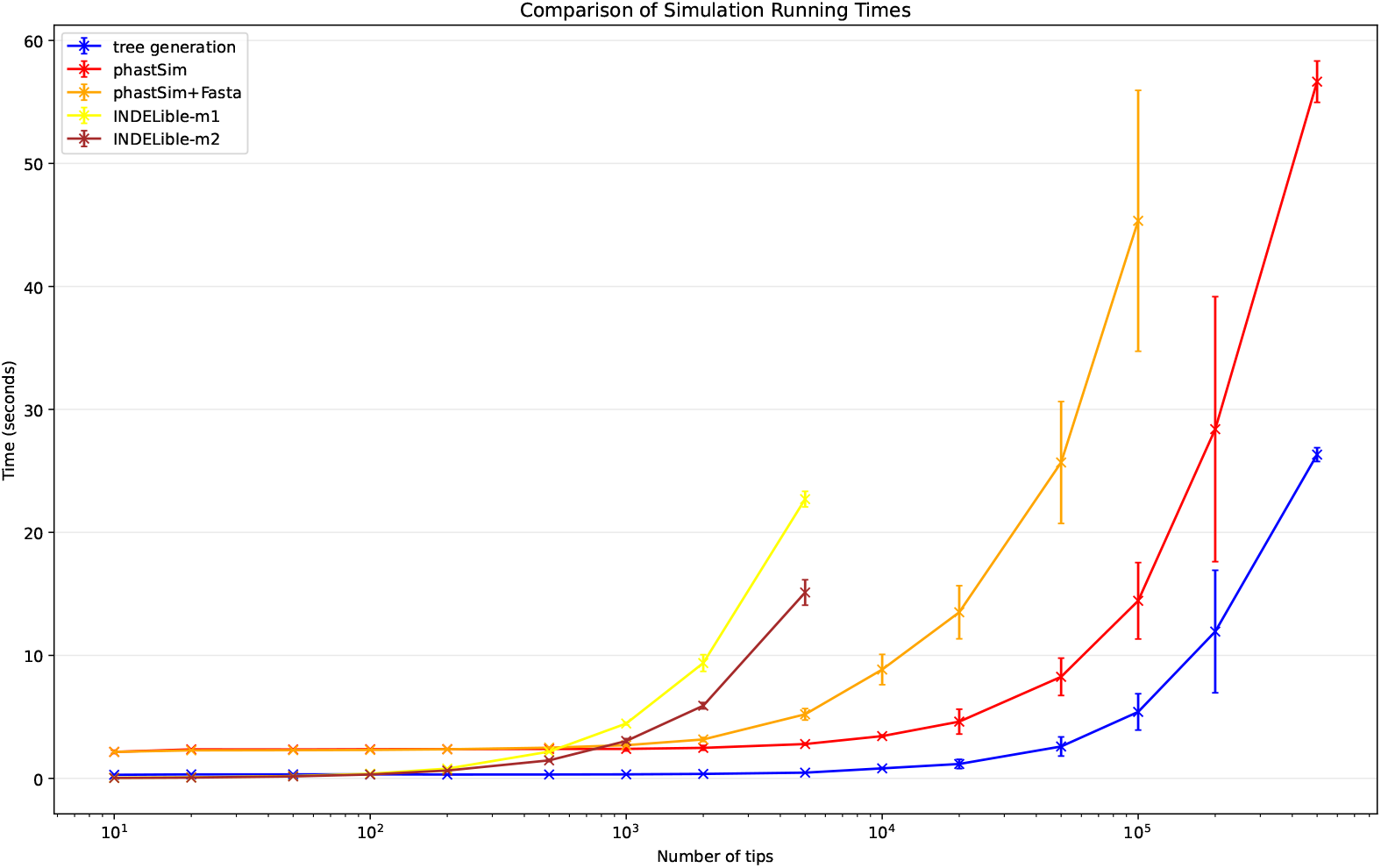
Comparison of running times of Indelible and phastSim simulators in a SARS-CoV-2 scenario with indels. In this scenario we compare phastSim against Indelbile-m1 and Indelible-m2 (the only other methods considered here that model indels), with uniform insertion and deletion rate of 0.1 and with indel length distribution of Geo(0.5). Each boxplot represents ten replicates.

### The impact of branch lengths

Probably the main limiting factor in the applicability of the approach presented here are tree branch lengths. Since the demand of our approach is affected linearly by the number of mutation events, and as we scale up the length of the tree we need to simulate more mutation events, then the length of the phylogenetic branches will significantly affect the performance of our approach. We can see that, in the SARS-CoV-2 scenario, the impact is not strictly linear (Fig 7). This is because there are additional factors which contribute to phastSim demand in addition to the number of mutation events. For example, one also has to consider the time to initialize the genome tree, which is linear in genome size, as well as the time to read, initialize, and traverse the input phylogenetic tree, which are linear in the number of tips.

**Fig 7.**
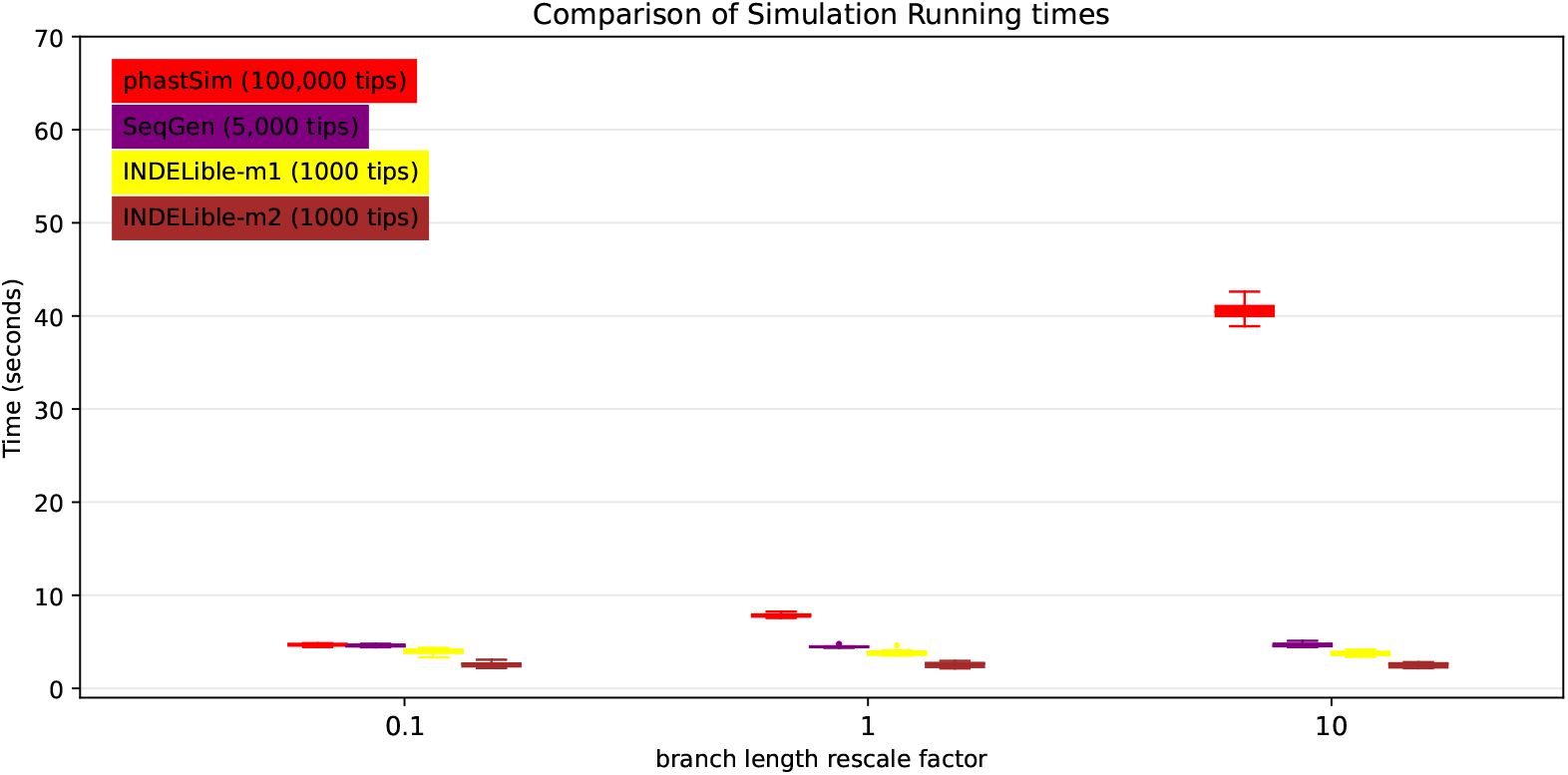
Comparison of running times of different simulators in a SARS-CoV-2 scenario after rescaling the tree branch lengths by different factors. On the Y axis we show the number of seconds it takes to perform simulations using different software. On the X axis is the rescaling factor we use to make the phylogenetic tree branch lengths longer or shorter. Colors are as in Fig 3. To aid the visual comparison, we use trees of different sizes for different simulators: 1000 tips for INDELible; 5,000 tips for Seq-Gen; 100,000 tips for phastSim.

Unexpectedly, the computational demands of Seq-Gen and INDELible-m1 seem not affected by the length of the branches. It is instead surprising to see that the computational demand of INDELible-m2 seems also not affected by the branch lengths, despite it using a Gillespie approach; the reason is that probably other factors, independent of the number of mutations, cause the bulk of the demand in this scenario.

## Discussion

We have introduced a new approach to simulating sequence evolution that is particularly efficient when used on phylogenies with many tips and with short branches. Our software phastSim implements this new algorithm and is implemented in Python, allowing it to be easily extended and combined with other Python packages. phastSim relies on the ETE 3 tree phylogenetic structure, and in particular it uses ETE 3 to read input phylogenetic trees. This allows flexibility in the phylogenetic tree input format. Furthermore, thanks to the fact that the efficiency of the algorithm is not affected by the complexity of the substitution model used, we allow a broad choice of evolutionary models, such as codon models with position-specific mutation rates and selective pressures. We also implement a new model of hypermutability to more realistically describe the mutational process in SARS-CoV-2. Also, we can efficiently simulate indel events, which are rarely modeled by other simulation packages.

We show that, compared with other simulators, phastSim is more efficient in the scenarios common to genomic epidemiology, that is, when simulating many closely related bacterial or viral genomes. Its particular efficiency with bacterial genomes means that it ideally matches the needs of software that simulate bacterial ancestral recombination graphs (e.g. [9, 41]). phastSim can also be easily run using the output of phylogenetic simulator, most relevantly VGsim [42] which allows fast simulations of very large and short phylogenies typical of SARS-CoV-2 and other genomic epidemiological scenarios, and which also allows the simulation of the effects of selection on the phylogenetic tree shape. phastSim is implemented as a Python package, which allows for easy integration into other Python pipelines.

In the future, it would be possible, and of interest, to expand the features of phastSim, in particular allowing a broader spectrum of models, for example allowing column-specific amino acid fitness profiles; also, it could be possible to implement the described algorithm in more efficient programming languages.

In conclusion, we have presented a novel algorithm, and corresponding software implementation phastSim, to efficiently simulate sequence evolution along large trees of closely related sequences. This new approach considerably outperforms other methods in the scenarios of genomic epidemiology, for example when simulating SARS-CoV-2 genome sequence datasets. This approach also allows for more realistic models of sequence evolution, allowing more efficient and accurate sequence data simulation and inference.

## Acknowledgments

We are very thankful to Vladimir Shchur for the valuable suggestions on our work.

## Code Availability

The code and data used for this project (except for the SARS-CoV-2 phylogenetic tree which falls under the restrictions of the GISAID terms of use) are available at https://github.com/NicolaDM/phastSim. phastSim can be easily installed across most platforms (see PyPI repository https://pypi.org/project/phastSim/) using the pip installer.

